# High-quality genome assemblies of four members of the *Podospora anserina* species complex

**DOI:** 10.1101/2023.10.24.563784

**Authors:** S. Lorena Ament-Velásquez, Aaron A. Vogan, Ola Wallerman, Fanny Hartmann, Valérie Gautier, Philippe Silar, Tatiana Giraud, Hanna Johannesson

## Abstract

The filamentous fungus *Podospora anserina* is a model organism used extensively in the study of molecular biology, senescence, prion biology, meiotic drive, mating-type chromosome evolution, and plant biomass degradation. It has recently been established that *P. anserina* is a member of a complex of seven, closely related species. In addition to *P. anserina*, high-quality genomic resources are available for two of these taxa. Here we provide chromosome-level annotated assemblies of the four remaining species of the complex, as well as a comprehensive dataset of annotated assemblies from a total of 28 *Podospora* genomes. We find that all seven species have genomes of around 35 Mbp arranged in seven chromosomes that are mostly collinear and less than 2% divergent from each other at genic regions. We further attempt to resolve their phylogenetic relationships, finding significant levels of phylogenetic conflict as expected from a rapid and recent diversification.

**Significance:** Here we provide a dataset of 28 annotated genomes from the *P. anserina* species complex, including chromosome-level assemblies of four species that lacked a reference genome. With this dataset in hand, biologists can take advantage of the molecular tools available for *P. anserina* to study evolutionary dynamics at the interphase between micro- and macroevolution, with particular emphasis on trait evolution, genome architecture, and speciation.

## Introduction

The filamentous fungus *Podospora anserina* (Order *Sordariales*) holds significant importance as a model for understanding ascomycete biology and beyond (Silar, 2013). It has proved particularly valuable in advancing the study of molecular biology, senescence, heterokaryon incompatibility, sexual reproduction, prion biology, meiotic drive, and plant biomass degradation (Grognet et al., 2014; Hamann and Osiewacz, 2018; Hartmann et al., 2021; Pinan-Lucarré et al., 2007; Silar, 2013, 2020; Vogan et al., 2022). Its reference genome was published as early as 2008 (Espagne et al., 2008), followed by chromosome-level assemblies of several wild-type strains (Vogan et al., 2021, 2019) and short-read population genomic data from Wageningen, the Netherlands, as well as a few strains from France and other localities (Ament-Velásquez et al., 2022). However, knowledge of its diversity, geographic distribution, ecology, and evolution lags behind. It is generally agreed that *P. anserina* is an obligately sexual coprophilous fungus, but there are observations of potential asexual spores (Boucher et al., 2017; Silar, 2020) and endophytic stages (Matasyoh et al., 2011). The name *P. anserina* itself has been riddled with taxonomic uncertainties (Ament-Velásquez et al., 2020; Silar, 2020), leading to confusion regarding the exact identity of the fungal material used in some studies. Unsurprisingly, a phylogenetic survey showed that many strains commonly regarded as *P. anserina* actually belong to at least six additional species scattered around the world (Boucher et al., 2017). Representatives of all these species have been sequenced with short-read technology, which was useful to explore the dynamics of recombination suppression around the mating-type locus (Hartmann et al., 2021). However, long-read data is necessary to understand the evolution and genetic basis of many traits. For example, the development of high-quality genomic resources of two of these species, *Podospora comata* and *Podospora pauciseta*, already provided important insights into the evolutionary dynamics of selfish genetic elements and genome architecture (Silar et al., 2018; Vogan et al., 2021, 2019).

As originally defined, only one or two strains are known for most members of the *P. anserina* species complex (Boucher et al., 2017), many available at the Westerdijk Fungal Biodiversity Institute Collection (identified with CBS numbers). All species have a similar morphology, mating system, and coprophilous habit, with the exception of the only known strain of *Podospora pseudocomata*, which was isolated from soil (Boucher et al., 2017; Hartmann et al., 2021). Despite their similarities, they are considered biological species, since there is reproductive isolation in the form of low mating success and female sterility in the hybrids (Boucher et al., 2017). Moreover, they are identifiable by differences at the fungal barcode ITS, as well as other nuclear markers (Boucher et al., 2017). Previous genomic comparisons showed that *P. anserina,P. comata*, and *P. pauciseta* are more than 98% identical in genic regions (Vogan et al., 2019), confirming that they are very closely related. However, their exact relationships remain unresolved. In this study, we generated chromosome-level annotated genome assemblies of the four remaining species (*Podospora bellae-mahoneyi, Podospora pseudoanserina, Podospora pseudopauciseta*, and *P. pseudocomata*), as well as short-read data from additional strains. In addition, we conducted a phylogenomic analysis to provide an evolutionary framework for addressing the variety of questions for which *Podospora* is well suited.

## Results and Discussion

### Genome assemblies and annotation

We isolated haploid cultures (of mating type + or -) from dikaryotic strains. From those, we selected one strain of each of the four species that lack a reference genome (hereafter, the focal strains) for Oxford Nanopore MinION and Illumina HiSeq sequencing (**Supplementary Table 1**). In addition, we sequenced with Illumina HiSeq a known strain of *P. pauciseta* (CBS 451.62+), the type strain of *P. pseudoanserina* (CBS 253.71+), and two newly collected *P. comata* strains (Wageningen Collection numbers Wa132+ and Wa133-) (**Supplementary Table 1**). Whole genome assemblies of Oxford Nanopore MinION data from the focal strains recovered mostly chromosome-level scaffolds that are highly collinear with the reference genome of *P. anserina* (**Figure 1**), although *P. pseudocomata* (strain CBS415.72-) has slightly more rearrangements. Thus, all species in the complex likely have seven chromosomes, a similar genome size of around 35 Mbp, and a repeat content ranging from more than 3% (*P. comata*) to around 7% (*P. pseudopauciseta*) (**Supplementary Table 1**) that is mostly concentrated in clusters (**Figure 1**). All assemblies gave comparable BUSCO numbers to the reference genome of *P. anserina* (**Supplementary Table 1**). Likewise, genome annotation using previously published RNAseq data of *P. anserina* and *P. comata* (Lelandais et al., 2022; Vogan et al., 2021) resulted in similar protein-coding genes numbers for assemblies produced with long-read data, from 11033 (*P. bellae-mahoneyi* strain CBS112042+) to 11727 (*P. anserina* strain T_G_+) genes, while the *P. anserina* reference itself has 10803 predicted protein-coding genes (**Supplementary Table 1**). The discrepancy in gene numbers with the reference is likely due to differences in the annotation method (with our pipeline, we recovered 11660 genes in the *P. anserina* reference assembly).

**Figure 1.**
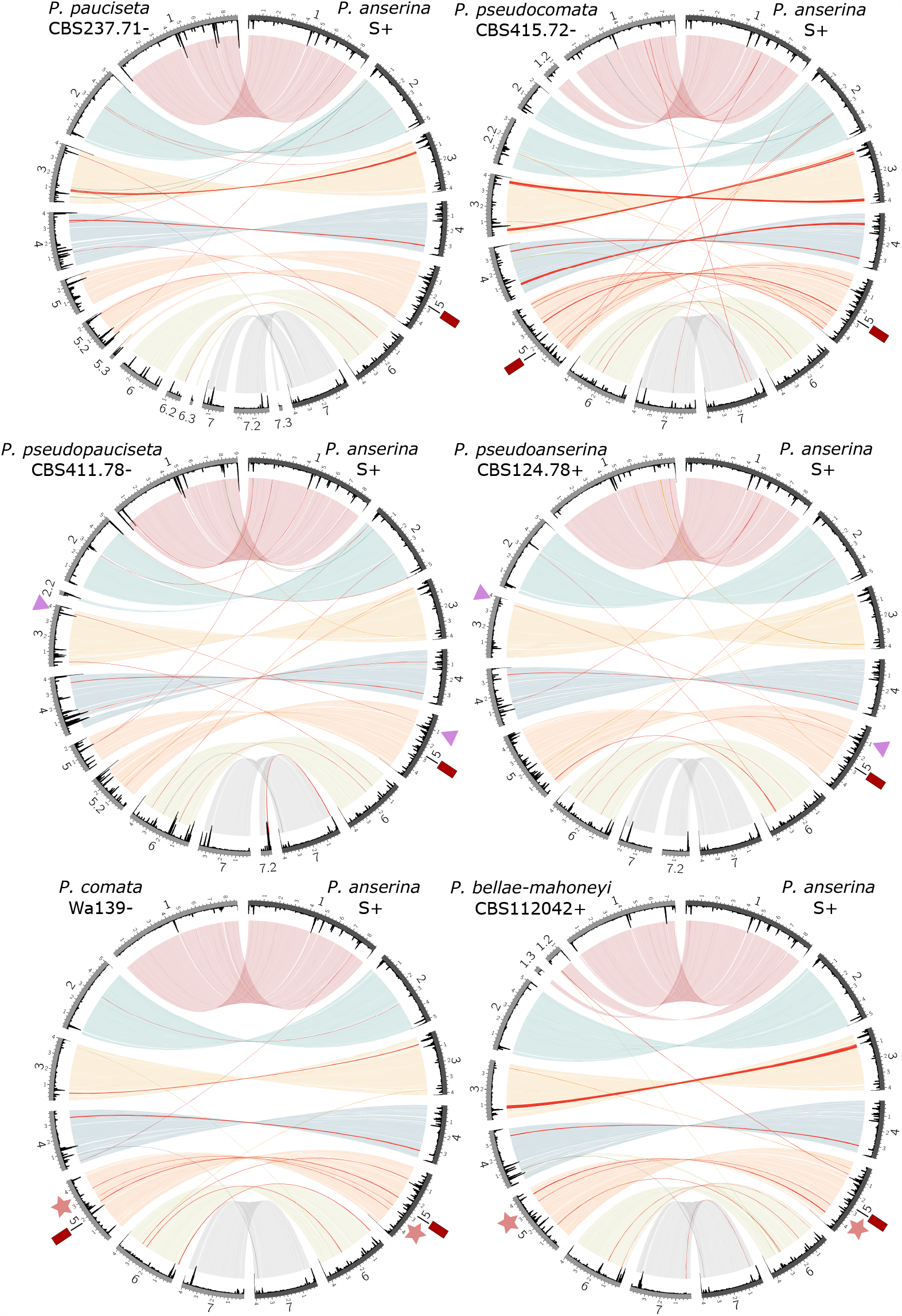
Circos plots comparing the reference assembly of *P. anserina* (strain S+, right side of each plot) to the best genome assembly of each of the other members of the species complex (left side). Light colors correspond NUCmer alignments (larger than 5kb) of the different chromosomes as defined in *P. anserina* (chr. 1: red, chr. 2: turquoise, chr. 3: yellow, chr. 4: blue, chr. 5: orange, chr. 6: olive green, chr 7: gray). Red links mark chromosome inversions or inverted translocations. The internal track in black is a histogram of repetitive element abundance calculated in sliding windows of 50 kb with steps of 10 kb. The stars and triangles mark shared structural variants (relative to *P. anserina*). The location of the insertion in chr. 5 discussed in the text is marked with a red square.

### Phylogenomics and comparative genomics

We aimed at providing a phylogenetic context of the *P. anserina* species complex by using all the available genomic resources (**Figure 2**; **Supplementary Table 1**). The closest known relative of the *P. anserina* species complex is *Cercophora samala*, strain CBS 307.81 (Ament-Velásquez et al., 2020). Preliminary phylogenomic analyses using CBS 307.81 as the outgroup placed the clade of *P. anserina* and *P. pauciseta* as sister to the other *Podospora* species (**Supplementary Figure 1**). However, this *C. samala* strain is in fact too divergent (around 86% identity in nuclear genic regions to any *Podospora* species) relative to the species complex (>98% identical to each other), potentially creating long-branch attraction (Emms and Kelly, 2017; Felsenstein, 1981). Hence, only *Podospora* strains were considered below, and we tentatively rooted the phylogeny using *P. anserina* and *P. pauciseta*.

**Figure 2.**
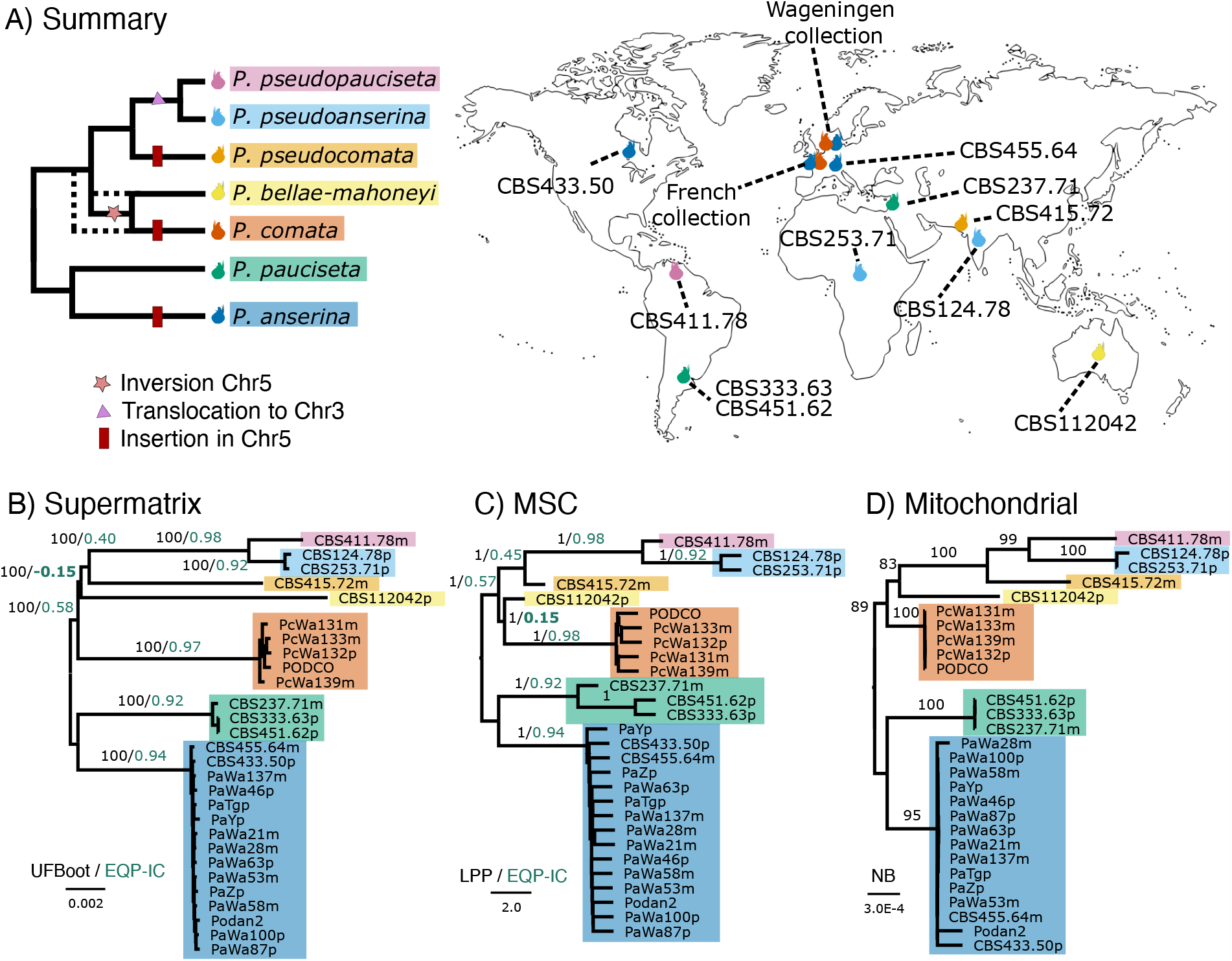
Phylogenetic relationships of the *Podospora* strains with genomic resources and their geographic distribution. (A) Summary cladogram based on the phylogenomic analyses and the detected structural variants, with dotted branches illustrating an alternative topology. Colored fruiting body cartoons mark the country where the different strains were sampled. Phylogenetic relationships were inferred from a supermatrix maximum-likelihood analysis (B) or a multispecies coalescent analysis (C) of nuclear genes, as well as a maximum-likelihood analysis of concatenated mitochondrial genes (D). Rooting is tentative based on analyses with *C. samala* as an outgroup. Branch lengths of the phylograms are drawn to scale as indicated by the scale bar (nucleotide substitutions per site in B and D and coalescent units in C). Different support metrics are shown next to their corresponding branches (within-species values are removed for clarity). The EQP-IC value of the conflicting branch is highlighted in bold. MSC: multispecies coalescent; UFBoot: ultrafast bootstraps; EQP-IC: extended quadripartition internode certainty; LPP: local posterior probability; NB: nonparametric bootstrap.

A summary of our results is found in **Figure 2A**. We inferred groups of single-copy orthologous (SCO) genes and used them to produce three different phylogenetic analyses (**Figure 2B-D**). All phylogenies resulted in well-supported species-level clades separated by very short internal branches, suggesting rapid diversification. Most relationships were congruent amongst analyses, except with regard to the relative positions of *P. comata* and *P. bellae-mahoneyi*. In the first analysis, a maximum-likelihood (ML) phylogeny produced from a supermatrix of 1000 nuclear SCO genes, *P. bellae-mahoneyi* is inferred as sister to the clade containing *P. pseudopauciseta, P. pseudoanserina*, and *P. pseudocomata* (**Figure 2B**). By contrast, a multispecies coalescent (MSC) analysis of all the 8596 SCO genes recovered *P. comata* and *P. bellae-mahoneyi* as sister taxa (**Figure 2C**). Lastly, a ML phylogeny of eight mitochondrial genes was in support of the nuclear supermatrix tree, albeit with modest bootstrap values (**Figure 2D**). To further explore this phylogenetic conflict, we obtained extended quadripartition internode certainty (EQP-IC) values (Zhou et al., 2020) for the two competing topologies. The EQP-IC score can range from 1 to -1. A score of 1 implies that all the SCO genes trees agree with a given branch, while a value of -1 reveals that all trees support an alternative topology. The score approaches 0 if the alternative topologies have similar frequencies amongst the gene trees (Zhou et al., 2020). As expected, we found intermediate positive values in most internal branches, which might be explained by incomplete lineage sorting (ILS) or introgression. The contesting branches in particular obtained values very close to 0, but there is slight support for the existence of a clade containing *P. comata* and *P. bellae-mahoneyi* (**Figures 2B and 2C**).

Taking advantage of the extensive collinearity between genomes, we also attempted to find phylogenetically informative structural variants. In support of the MSC analysis, we found a medium-scale (33.89 kb, 13 genes) inversion in chromosome 5, relative to *P. anserina*’s genome, that is shared between *P. comata* and P. *bellae-mahoneyi* (**Figures 1 and 2A**). In addition, we detected a shared translocation from chromosome 5 to chromosome 3 between *P. pseudoanserina* and *P. pseudopauciseta*, supporting their close sister-relationship already observed in the phylogenetic analyses (**Figures 1 and 2A**). However, in contradiction with our phylogenies, we found a region in chromosome 5 that is only present in *P. anserina, P. comata*, and *P. pseudocomata* (**Figures 1 and 2A**). This region ranges from over 40 kb in *P. anserina* to just over 6 kb in P. *pseudocomata* and contains a number of genes and different transposable elements (TEs). Upon closer inspection, we found that the edges of this region are flanked by directed repeats of 4 bp, suggesting that this region represents a TE-mediated insertion, resembling the behavior of large TEs found in Pezizomycetes, including *Podospora* (Gluck-Thaler et al., 2022; Vogan et al., 2021). Its phylogenetic distribution might also be a consequence of ILS or introgression.

## Conclusion

Here, we present high-quality annotated genome assemblies of four members of the *P. anserina* species complex. Together with the already available genomic resources, this new data builds an evolutionary framework for further in-depth studies of this group of fungi. We provide a general idea of the genomic architecture and relationships between the sampled *Podospora* lineages, while illustrating the high levels of phylogenetic conflict that are typical of rapid species radiations. Moreover, we make available a comprehensive genomic dataset of 28 strains for the study of fungal biology and evolution at shallow divergence scales. Combined with the wealth of molecular biology tools available for the model species *P. anserina*, this dataset can be used to explore the evolution and function of pangenome content including metabolic clusters, selfish genetic elements like meiotic drivers and transposable elements, as well as the buildup of reproductive barriers in filamentous fungi.

## Materials and Methods

All bioinformatic pipelines were done in Snakemake v. 7.25.0 or v. 7.32.3 (Mölder et al., 2021), and are available at https://github.com/SLAment/PodosporaGenomes, unless otherwise stated. Some of these pipelines rely on the Environment for Tree Exploration (ETE3) toolkit v. 3.1.3 (Huerta-Cepas et al. 2016).

### Fungal material

For detailed information about the strains used, see **Supplementary Table 1**. The strains with code starting with “CBS” were originally obtained from the CBS-KNAW Collection (https://wi.knaw.nl/Collection). In addition, we isolated two new strains of *P. comata* from rabbit dung collected in the area between Wageningen and Arhem, the Netherlands (51°58’41.8”N 5°50’39.6”E, locality Unksepad Oosterbeek) in September of 2016, which were deposited in the Wageningen Collection at the Laboratory of Genetics of the Wageningen University and Research (codes Wa132 and Wa133). These *P. comata* strains were obtained by incubating the rabbit dung pieces on 9 cm Petri dishes at room temperature until *Podospora*-like perithecia developed (around 10 days). The perithecia were identified based on gross morphology, and moved from the dung into a water agar plate covered with an NC 45 membrane filter (Schleicher & Schuell, Dassel, Germany) and pierced with a sterilized needle to retrieve an ascus with 4 spores. The four spores were isolated into germination media (PASM2 with 5g/L ammonium acetate; Vogan et al., 2019). Two days after germination, the mycelium was stored by inoculating PASM0.2 plates (Vogan et al., 2019). Only one spore was used as representative culture, but all siblings were stored in the Wageningen Collection. For all *Podospora* strains, we obtained haploid cultures by letting them undergo selfing and retrieving sexual ascospores with a single haploid nucleus (i.e., a monokaryotic spore). We indicate the mating type of the monokaryotic cultures with either a + or a - next to the strain number. The sequenced *C. samala* strain was a monokaryotic F1 progeny from strain CBS307.81 (of - mating type, hence referred to as CBS307.81-) (see Hartmann et al., 2021). Notice that a total of 106 *P. anserina* strains from Wageningen were sequenced previously (Ament-Velásquez et al., 2022), but only those with long-read data were selected as representatives for our analyses. All strains from outside of Wageningen or from the other species with available sequence data were also included (**Supplementary Table 1**).

### DNA extraction and sequencing

For Illumina sequencing, we grew the haploid strains on Petri dishes of either M2 medium (CBS124.78+ and CBS307.81-) or PASM0.2 (other strains) for 3 to 4 days (Silar, 2020; Vogan et al., 2019). We scraped mycelium off the plates in order to obtain about 80-200 mg of mycelium per strain and stored it in 1.5 mL Eppendorf tube at -80°C for at least 24 hours before extraction. For most strains, whole genome DNA was extracted with the Fungal/Bacterial Microprep kit (Zymo; https://zymoresearch.eu/). In the case of CBS124.78+ and CBS307.81-, the mycelium was lyophilized for about 20 hours, and DNA was extracted using the commercial Nucleospin Soil kit from Macherey Nagel. Paired-end libraries (150 bp reads) were sequenced by either the SNP and SEQ Technology platform (SciLifeLab, Uppsala, Sweden) on the Illumina HiSeq X platform (most strains), or by the high throughput sequencing core facility of I2BC, Université Paris-Saclay (Centre de Recherche de Gif – http://www.i2bc.paris-saclay.fr/) on the Illumina NextSeq500 platform (CBS124.78+ and CBS307.81-).

For MinION Oxford Nanopore sequencing, we grew the strains in liquid cultures of 3% malt extract solution as in Vogan et al. (2019). High molecular-weight DNA was extracted as in Sun et al. (2017), using the Genomic Tip G-500 columns (Qiagen) and the PowerClean DNA Clean-Up kit (MoBio Labs). The strain CBS 411.78-was prepared and sequenced with the ligation kit SQK108 in a one-pot reaction using 500 ng DNA for end prep and ligation (NEB Ultra-II ligase) and sequenced on an R9.4.1 flowcell. In addition, a rapid barcoding (RBK004) was made for CBS 411.78-to get sufficient coverage for assembly. Similarly, CBS 415.72-was also sequenced using both the RBK004 and LSK108 kit on an R9.4.1 flowcell to maximize yield. CBS 112042+ and CBS 124.78+ were sequenced on R9.4.1 flowcells using the LSK108 kit with 3 μg DNA as input for the end prep reaction (NEB ULTRA-II EP, 20 min. at 20°C and 20 min. at 65°C). A bead purification (SpeedBeads, GE) was done before ligation to deplete short fragments and 1.5 μg DNA per sample was ligated to 20 μl AMX 1D using Blunt/TA ligase (30 min.).

### Genome assembly

The paired-end HiSeq Illumina reads were cleaned from adapters using cutadapt v. 1.13 (Martin 2011) and Trimmomatic v. 0.36 (Bolger, Lohse, and Usadel 2014) with the options: ILLUMINACLIP:adapters.fasta:1:30:9 LEADING:20 TRAILING:20 SLIDINGWINDOW:4:20 MINLEN:30, as in Vogan et al. (2019). We used both forward and reverse paired-end reads in downstream analyses. *De novo* assemblies were produced as in Vogan et al. (2019). Specifically, we assembled the MinION reads with mean Phred quality (QV) above 9 and longer than 1kb using Minimap2 v. 2.11 and Miniasm v. 0.2 (H. Li 2018, 2016). Racon v. 1.3.1 (Vaser et al. 2017) was used twice to polish the resulting assemblies based on the unfiltered reads. We further polished five times using the Illumina reads with Pilon v. 1.22 (Walker et al. 2014), which were mapped using BWA v. 0.7.17 (H. Li and Durbin 2010), with PCR duplicates marked by Picard v. 2.18.11 (http://broadinstitute.github.io/picard/), and with local indel re-alignment from the Genome Analysis Toolkit (GATK) v. 3.7 (Van der Auwera et al. 2013). For the strains without long-read data, we ran SPAdes v. 3.12.0 (Bankevich et al. 2012) with the k-mers 21,33,55,77 (Wa131-, Wa132+, Wa133-, Wa139-, CBS333.63+, CBS451.62+, CBS253.71+) or 21,29,37,45,53,61,79,87 (CBS307.81-) and the *--careful* option. Due to high numbers of read pairs with read mates mapping on different chromosomes during the read mapping procedure (about 20% of mapping read pairs; analyses not shown), we used each paired-end sequencing run as two independent single-end sequencing runs for read mapping and de novo assembly of CBS124.78+ and CBS307.81-, respectively (see Hartmann et al., 2021).

In the case of samples with long-read data, the scaffolds were assigned to chromosomes and re-oriented by mapping them to the reference genome of the strain S (Espagne et al. 2008), which is available at the Joint Genome Institute MycoCosm website (https://mycocosm.jgi.doe.gov/Podan2/Podan2.home.html) as “Podan2”. Mapping was performed with MUMmer package v. 4.0.0beta2 (Kurtz et al. 2004) using the parameters *-b 2000 -c 200 --maxmatch*. Contigs smaller than 100 kb that contained rDNA repeats or mitochondrial sequences were discarded (except for the largest mitochondrial contig). Genome quality statistics were calculated with QUAST v. 4.6.3 (Mikheenko et al. 2016). Mean depth of coverage was obtained using QualiMap v.2.2 (Okonechnikov, Conesa, and García-Alcalde 2016). We also used BUSCO v. 5.3.1 (Manni et al., 2021) with the 3817 Sordariomycetes_odb10 ortholog set to assess assembly completeness. As dependencies we used BLAST suit 2.12.0+ (Camacho et al., 2009), AUGUSTUS v. 3.4.0 (Stanke and Waack, 2003), and HMMER v. 3.2.1 (Mistry et al., 2013). In addition, we included in our analyses the previously produced assemblies of *P. anserina* (Espagne et al., 2008; Vogan et al., 2021, 2019), *P. comata* (Silar et al., 2018; Vogan et al., 2021) and *P. pauciseta* (Vogan et al., 2019).

We verified the correct assembly of the mitochondrial contig in the focal strains CBS 112042+, CBS 124.78+, CBS 411.78- and CBS 415.72-by visual inspection of long- and short-reads mapping. We found a misassembly in the contig of CBS 415.72-around the first exon of the *cox1* gene. Hence, we extracted the long reads mapped to this mitochondrial contig using the *bam2fq* option of SAMtools v. 1.17 (Danecek et al., 2021) and reassembled them with Flye v. 2.9.1 (Kolmogorov et al., 2020) with the arguments *--iterations 2 --meta --keep-haplotypes*. We recovered two circular contigs, one of which proved to be formed by multiple tandem repeats of the first part of *cox1*, a configuration known as α senDNA or plDNA (Cummings et al., 1985; Hamann and Osiewacz, 2018). The other contig corresponded to the full mitochondrion. We polished the mitochondrial contig three times using the Illumina reads as above and discarded the plDNA. We manually re-circularized the mitochondrial contigs of the focal strains to avoid breaking genes.

### Genome annotation

The annotation of all genomes was done with a modified version of a previous pipeline (Vogan et al., 2021). Briefly, we used previously produced (Vogan et al., 2019) training files for SNAP release 2013-11-29 (Lomsadze et al., 2005) and GeneMark-ES v. 4.38 (Lomsadze et al., 2005; Ter-Hovhannisyan et al., 2008) within the program MAKER v. 3.01.04 (Holt and Yandell 2011; Campbell et al. 2014) to generate gene models for all species. MAKER was run with the following dependencies: BLAST suit 2.13.0+, tRNAscan-SE v. 1.3.1 (Lowe and Eddy, 1997), Exonerate v. 2.4.0 (Slater and Birney, 2005), and RepeatMasker v. 4.1.0 (http://www.repeatmasker.org/). As external protein evidence we used the sequences from the PODANS_v2016 annotation (Lelandais et al., 2022) and from the reference genome of *P. comata* (PODCO; Silar et al., 2018), along with a small set of manually curated proteins (available in the GitHub repository). We also used transcript models as external evidence from two sources: the curated set of mRNA with defined transcription starts and ends from PODANS_v2016, and transcript models of published RNAseq data of *P. anserina* and *P. comata* (Vogan et al., 2021, 2019). The latter were produced as in Vogan et al. (2021), using STAR v. 2.7.10b (Dobin et al., 2013), Cufflinks v. 2.2.1 (Trapnell et al., 2010), and TransDecoder v. 5.7.0 (Haas et al., 2013). Additionally, we used the custom library “PodoTE-1.00” (available at https://github.com/johannessonlab/SpokBlockPaper/blob/master/Annotation/data/) to annotate repeated elements in all species (Vogan et al., 2021). Manual curation of the output of RepeatModeler v. 1.0.8 (http://www.repeatmasker.org/RepeatModeler/) from long-read assemblies, as produced by the pipelines *PaTEs.smk* and *TEManualCuration.smk* (https://github.com/johannessonlab/SpokBlockPaper), revealed no new TEs in the *P. anserina* complex except for two elements in *P. pseudocomata*: a novel LINE in that we named *kermit* (no other LINE elements have been identified in the other *Podospora* species), and a new *Ty3* element related to *Yeti* (Hamann et al. 2000) that we called *sasquatch*. In addition, we found that the previously unclassified element *leptodactylodon* is an LTR element and that *P. pseudocomata* has a full copy of the *rana* LTR, which was previously only known from solo elements (Espagne et al., 2008). These new sequences are part of PodoTE-1.00 library. The total repeat content in a genome was estimated from the output of RepeatMasker as the percentage of sites annotated as repeats out of the total in the assembly (excluding mitochondrial contigs) with the script “totalcovergff.py” v. 2.2 available at https://github.com/SLAment/Genomics/tree/master/GenomeAnnotation.

Functional annotation was done with Funannotate v. 1.8.15 (Palmer & Stajich, 2020) using the *annotate* function with the dependencies HMMER 3.3.2 (Mistry et al., 2013), Diamond v. 2.1.6 (Buchfink et al., 2021) with the UniProt DB version 2023_01, InterProScan v. 5.62-94.0 (Blum et al., 2021; Jones et al., 2014), bedtools v. 2.30.0 (Quinlan and Hall, 2010), Eggnog-mapper v. 2.1.10 (Cantalapiedra et al., 2021) with the database emapperdb-5.0.2, and the fungal version of antiSMASH v. 6.1.1 (Blin et al., 2021). The input assemblies were soft-masked with RepeatModeler externally with the PodoTE-1.00 library as above.

The original gene names of the reference S+ strain genome (Espagne et al., 2008; Lelandais et al., 2022) are extensively used by the *Podospora* scientific community and follow a convention that marks the chromosome name and the gene number within the chromosome. For example, “Pa_1_3060” is the gene 3060 in chromosome 1. Hence, we took advantage of the high collinearity between the *Podospora* genomes to identify one-to-one orthologs using BLASTn and name them accordingly. If a focal gene had a single, high-quality hit in the PODANS_v2016 annotation (Lelandais et al., 2022), and this hit was itself the only (high quality) hit of the focal gene, these were considered one-to-one orthologs. A high quality hit had maximum e-value of 0.001 and an identity equal or higher than 98% (for *P. anserina* strains), 93% (*P. comata* or *P. pauciseta*), 90% (the other *Podospora* species) or 70% (*C. samala*). This simple strategy assigned more than 90% of genes to one-to-one orthologs in all strains. The IDs of these genes were then set to reflect their relationships to the reference genome. For example, the one-to-one ortholog of “Pa_1_3060” was named “QC762_103060” in the strain CBS415.72-, where QC762 is the strain’s NCBI locus tag.

Mitochondrial annotation was done with the online version of MFannot (Lang et al., 2023), setting the genetic code to 4 (https://megasun.bch.umontreal.ca/apps/mfannot/; consulted during the second half of July 2023). The resulting tbl format was modified with the script *MFannot4ncbi.py* to make it closer to NCBI requirements, and further transformed to a gff3 file with the script *tbl2gff.py* (available at https://github.com/SLAment/Genomics/blob/master/GenomeAnnotation). We performed manual curation of all the canonical protein coding genes, as well as the ribosomal large and small subunit genes, to match the *P. anserina* reference (Genbank accession number NC_001329.3). Considering the high Nanopore error rate and the poor mapping of Illumina reads to mitochondrial contigs during polishing of CBS 112042+ and CBS 411.78-(due to the presence of multiple mitochondrial contigs), we verified that the genes sequence matched the assembly of short reads and corrected them when necessary. The final gff3 file was transformed back to tbl format with *gff3TOtbl.py* (also in the repository above) for submission to NCBI.

### Comparative genomics

In order to explore collinearity among the *Podospora* species, we used the NUCmer program of the MUMmer package v. 4.0.0beta2 (Kurtz et al., 2004) with parameters *-b 2000 -c 2000 -- maxmatch* to align the long-read assembly of a representative of each species (CBS 237.71-, Wa139-, CBS 112042+, CBS 415.72-, CBS 411.78-, and CBS 124.78+) against Podan2. We further calculated the coverage distribution of repetitive elements along chromosomes using the RepeatMasker annotation and by dividing the genome in windows of 50 kb with steps of 10 kb using the utilities *makewindows* and *coverage* of BEDtools v. 2.29.0 (Quinlan, 2014; Quinlan and Hall, 2010). Both the alignments and coverage distributions were plotted using Circos v.

0.69.6 (Krzywinski et al., 2009). We removed all alignments smaller than 5 Kb to exclude the most abundant transposable elements. All inversions and translocations detected in the NUCmer output were manually inspected to identify homologous events between species and marked in the final Circos figure. The shared inversion in *P. comata* and *P. bellae-mahoneyi* compared to the other species is located in the Podan2 genome at coordinates 3440773-3474671 in chromosome 5. We verified that the edges are identical. Similarly, the shared translocation from chromosome 5 in the other species to chromosome 3 in *P. pseudoanserina* and *P. pseudopauciseta* is located at Podan2 coordinates 966739-973648. The insertion shared by *P. anserina, P. comata* and *P. pseudocomata* is located at coordinates 2752403-2784937 in chromosome 5 of Podan2.

### Phylogenomic analyses

In order to resolve the relationships between the *Podospora* species, we inferred SCO groups by running OrthoFinder v. 2.5.2 (Emms and Kelly, 2019) with a single representative per species: strains S+ (Espagne et al., 2008; Lelandais et al., 2022), T_D_+ (Silar et al., 2018), CBS 237.71-(Vogan et al., 2019), and our focal strains CBS 124.78+, CBS 411.78-, CBS 415.72-, and CBS 112042+ (**Supplementary Table 1**).

OrthoFinder was run with the proteins predicted for the chosen strains. However, the low level of divergence within the species complex makes the proteins largely uninformative. Hence, once SCO groups were defined, we used the ortholog of the reference S+ as a BLASTn query to retrieve the nucleotide sequences of the corresponding homologs (including introns) in all the *Podospora* strains in our dataset (**Supplementary Table 1**). We kept only the orthogroups with a single sequence per strain, resulting in a total of 8596 SCO groups. These were then aligned with MAFFT v. 7.407 (Katoh and Toh, 2008) with the options *--adjustdirection --anysymbol -- maxiterate 1000 --retree 1 --localpair*. We inferred ML trees of each alignment using IQ-TREE v. 2.2.3 (Hoang et al., 2017; Nguyen et al., 2015) with parameters *-m MFP -seed 1234 -bnni -- keep-ident -bb 1000*. To reduce noise in our dataset, we collapsed branches with ultrafast bootstraps (UFBoots) support lower than 95% into polytomies using Newick utilities v. 1.6 (Junier & Zdobnov 2010). These trees were then given to ASTRAL v. 5.7.3 (Zhang et al. 2018) to construct a MSC phylogeny. In addition, we randomly selected 1000 of the SCO to form a concatenated alignment (supermatrix) of 1,732,364 sites (39,771 (2.4%) informative) and produced a ML phylogeny with IQ-TREE as above. In order to evaluate the level of conflict within the SCO trees with respect to both the supermatrix ML and MSC phylogenies, we calculated the Extended Quadripartition Internode Certainty (EQP-IC) score with the program QuartetScores v. 1.0 (Zhou et al., 2020) using the unrooted reference trees and the SCO trees with collapsed low-support branches from above as the evaluation set.

To complement the nuclear data, we extracted the coding sequence of seven mitochondrial genes with one or no introns (*atp6, cox2, cox3, nad2, nad3, nad4*, and *nad6*), as well as the small subunit ribosomal RNA gene (*rns*), from both the long- and short-read assemblies. As the polishing often failed for mitochondrial contigs, we gave priority to the short-read assembly sequences when discrepancies occurred. We excluded all introns since these are known to be polymorphic in at least *P. anserina* (Belcour et al., 1997). Moreover, the presence of introns and other regions of the mitochondrial genome can depend on the age of the mycelium (Cummings et al., 1985; Hamann and Osiewacz, 2018), and preliminary analyses with multiple-intron genes revealed that polymorphic sites are often correlated with intron presence/type rather than phylogenetic signal, in particular at the edges of exons. The final concatenated alignment of the eight genes contained 8655 sites, including 34 informative sites. We produced ML trees with IQ-TREE as above.

OrthoFinder can estimate the root in a set of species using patterns of gene duplications in the input proteomes (Emms and Kelly, 2017). In our analysis, this method put the root between a clade containing *P. anserina* (S+) and *P. comata* (T_D_+) and a sister clade containing all the other species. This result is at odds with all of our analyses above, where *P. anserina* and *P. pauciseta* are sister species. We found this result was driven by the shared gene content in the chromosome 5 insertion, as seven annotated genes within this region are exclusive to those two species and absent in the homologous region of *P. pseudocomata*. Additional manual attempts to find shared gene duplications across the complex failed. Hence, we did not consider this rooting method further.

## Data availability

The genome assemblies and raw sequencing data are available at GenBank under the BioProject PRJNA685103 (see **Supplementary Table 1**). The reads of the strains sequenced with long-read technology in this study have BioSamples accessions SAMN17076437 (CBS124.78+), SAMN17076438 (CBS411.78-), SAMN17076439 (CBS415.72-), and

SAMN17076440 (CBS112042+). The strains newly sequenced with Illumina have BioSamples with accessions SAMN37845441 (Wa132+), SAMN37845442 (Wa133-), SAMN37845443 (CBS 451.62+) and SAMN37845444 (CBS 253.71+). In addition, all genome assemblies, annotations in gff3 format, and alignments are available in Dryad Digital Repository (https://doi.org/10.5061/dryad.1vhhmgr0j). Scripts and snakemake pipelines are available at https://github.com/SLAment/PodosporaGenomes.

## Acknowledgments

This work was supported by the European Research Council (ERC) grants ERC-2014-CoG (project 648143 SpoKiGen) to H.J. and advanced grant 832352 EvolSexChrom, as well as the Louis D. Foundation (Institut de France) to T.G., a Marie Curie European grant (PRESTIGE-2016-4-0013), a Fondation des Treilles grant to F.E.H., and support from the Lars Hierta Memorial Foundation, The Nilsson-Ehle Endowments of the Royal Physiographic Society of Lund, and the Swedish Research Council (grant 2022-00341) to S.L.A.-V. We acknowledge the support given by the National Genomics Infrastructure (NGI) / Uppsala Genome center on massive parallel DNA sequencing. The computations were performed on resources provided by the National Academic Infrastructure for Supercomputing in Sweden (NAISS) and the Swedish National Infrastructure for Computing (SNIC) at Uppsala Multidisciplinary Center for Advanced Computational Science (UPPMAX) partially funded by the Swedish Research Council through grant agreements no. 2022-06725 and no. 2018-05973.

## Supplementary Materials

**Supplementary Figure 1.**
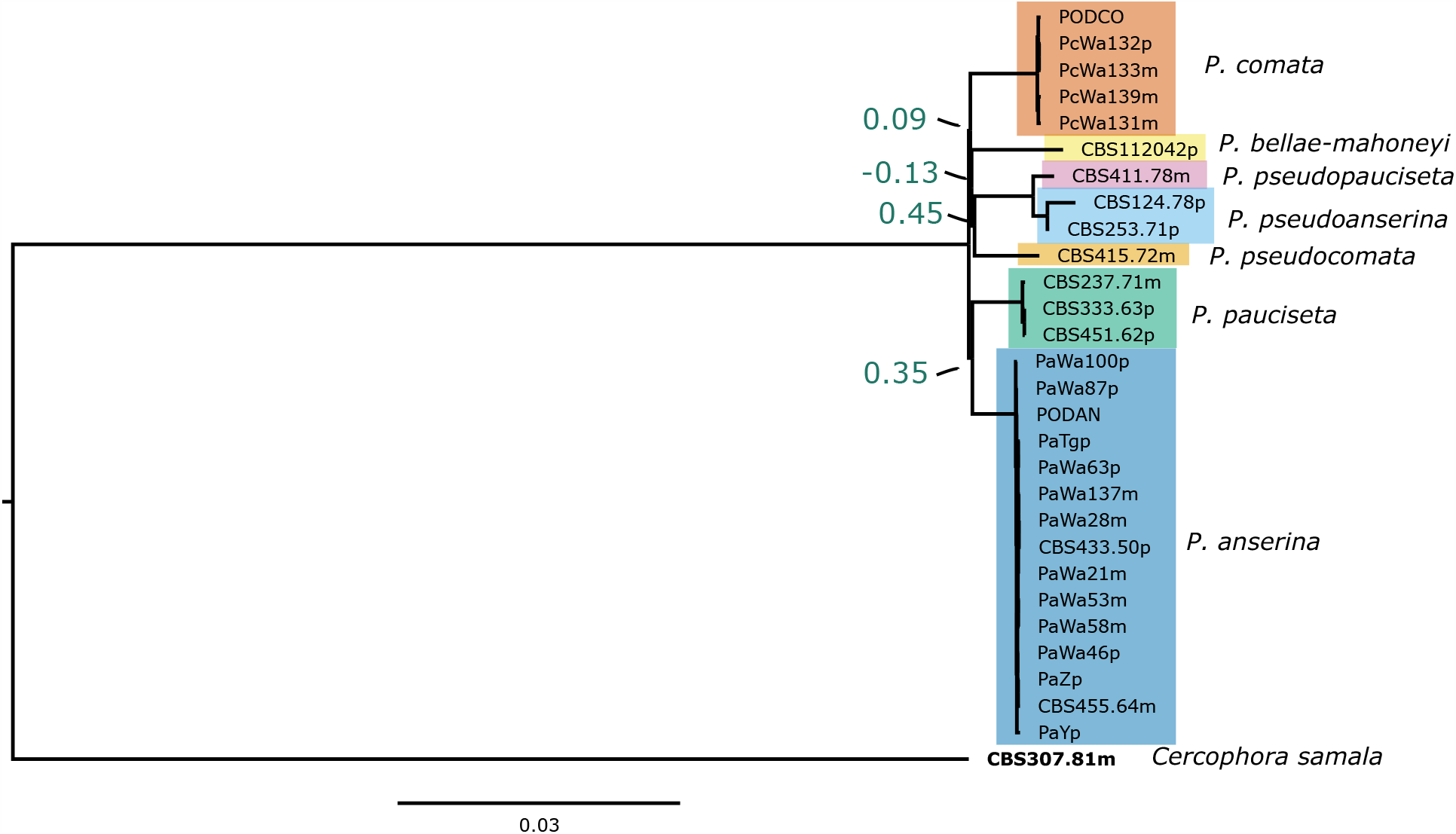
Maximum-likelihood analysis of 1000 concatenated nuclear genes. Branch lengths are drawn to scale as indicated by the scale bar (nucleotide substitutions per site). All species-level branches have ultrafast bootstraps (UFBoot) support of 100. Green scores correspond to extended quadripartition internode certainty (EQP-IC) values. The clade containing all *Podospora* species other than *P. anserina* and *P. pauciseta* has an EQP-IC of 0.09, a value much lower compared to the analysis without an outgroup, likely as a result of instability in the position of *C. samala* across gene phylogenies.

**Supplementary Table 1.** Metadata and genome assembly statistics of all the strains included in this study.

